# Pango lineage designation and assignment using SARS-CoV-2 spike gene nucleotide sequences

**DOI:** 10.1101/2021.08.10.455799

**Authors:** Áine O’Toole, Oliver G. Pybus, Michael E. Abram, Elizabeth J. Kelly, Andrew Rambaut

## Abstract

More than 2 million SARS-CoV-2 genome sequences have been generated and shared since the start of the COVID-19 pandemic and constitute a vital information source that informs outbreak control, disease surveillance, and public health policy. The Pango dynamic nomenclature is a popular system for classifying and naming genetically-distinct lineages of SARS-CoV-2, including variants of concern, and is based on the analysis of complete or near-complete virus genomes. However, for several reasons, nucleotide sequences may be generated that cover only the spike gene of SARS-CoV-2. It is therefore important to understand how much information about Pango lineage status is contained in spike-only nucleotide sequences. Here we explore how Pango lineages might be reliably designated and assigned to spike-only nucleotide sequences. We survey the genetic diversity of such sequences, and investigate the information they contain about Pango lineage status. Although many lineages, including the main variants of concern, can be identified clearly using spike-only sequences, some spike-only sequences are shared among tens or hundreds of Pango lineages. To facilitate the classification of SARS-CoV-2 lineages using subgenomic sequences we introduce the notion of designating such sequences to a “lineage set”, which represents the range of Pango lineages that are consistent with the observed mutations in a given spike sequence. These data provide a foundation for the development of software tools that can assign newly-generated spike nucleotide sequences to Pango lineage sets.

## Introduction

During the SARS-CoV-2 pandemic virus genome sequences have been generated and shared in unprecedented numbers. More than 2.2 million SARS-CoV-2 sequences have been deposited in the online database GISAID (Shu & McCauley 2017), as of 5^th^ June 2021. Analyses of virus genomes from the COVID-19 pandemic have revealed the international dissemination of the virus (e.g. du Plessis et al. 2021; Lemey et al. 2021), been used to support contact tracing and outbreak control (e.g. Li et al. 2021; Geoghegan et al. 2020), and enabled the discovery and surveillance of variants of concern (e.g. Rambaut et al. 2020; Faria et al. 2021; Tegally et al. 2021) or other lineages of virological or epidemiological interest.

Systems of nomenclature are common throughout biology. In virology they are used to organise and simplify observed virus genetic diversity at different taxonomic levels, and allow virologists, epidemiologists and public health officials to communicate precisely and unambiguously (e.g. Walker et al. 2019; Smith et al. 2014). In the case of the COVID-19 pandemic, at least four naming systems have been developed and used to identify and name genetically-distinct SARS-CoV-2 types and lineages (Alm et al. 2020; Konings et al. 2021). The Pango dynamic nomenclature system was developed and published in early 2020 (Rambaut et al. 2020; https://pango.network) and has since become a widely-used tool worldwide for SARS-CoV-2 classification. It differs from other SARS-CoV-2 nomenclature systems (such as GISAID and NextStrain) in that it (i) aims to capture the ongoing, leading edge of pandemic transmission, (ii) its lineages are intended to represent epidemiologically-relevant events, and (iii) it examines the phylogenetic structure of the pandemic at high resolution. The Pango nomenclature therefore contains a large number of lineages (currently >1,300; https://cov-lineages.org) covering the entire genetic diversity of SARS-CoV-2, some of which are genetically very similar to each other. In contrast, the ‘Greek letter’ system recently proposed by the WHO Virus Evolution Working Group (Konings et al. 2021) is intended for public communication purposes and provides labels only for a small number of variants of concern (VOCs) and variants of interest (VOIs). The WHO variant of concern labels Alpha, Beta, Gamma and Delta correspond to the Pango lineages B.1.1.7, B.1.351, P.1, and B.1.617.2, respectively.

The Pango nomenclature was designed to classify complete or near complete SARS-CoV-2 genomes. New Pango lineages are designated only if the lineage contains a sufficient number of sequences with high genome coverage. Specifically, a sequence is defined as high coverage if <5% of nucleotides sites across the whole genome (excluding UTRs) are missing or represented by IUPAC ambiguity codes (Rambaut et al. 2020). Although this criterion is required by Pango to “designate” a genome to a lineage, less complete genomes can be “assigned” to a lineage using the software tool *pangolin* (O’Toole et al. 2021). Thus sequence “designation” is a formal, definitive statement about lineage membership of a genome, whereas “assignation” is an estimate or inference of the lineage to which a sequence most likely belongs (https://pango.network). Tools for lineage assignation, such as the machine learning approach implemented in *pangolin* are important because it takes time for new sequences to be evaluated and designated by the Pango team, and because only a subset of of SARS-CoV-2 sequences available on GISAID meet the strict coverage criterion above.

One notable class of incomplete SARS-CoV-2 sequences comprise the complete nucleotide coding sequence of the spike glycoprotein. The spike protein is of particular biological importance because it is the primary target of natural and vaccine-elicited humoral immunity, it is the target of most therapeutic and prophylactic monoclonal antibodies, and it contains the virus’ receptor binding domain, which binds the human host cell receptor ACE2 with high affinity (e.g. Hu et al. 2021). Spike-only nucleotide sequences, rather than complete genomes, may be generated by researchers for several reasons. For example, many immunological investigations do not require information about the remainder of the virus genome, and a sub-genomic sequencing approach may sometimes generate higher quality spike sequences from high Ct samples. Further, some laboratories may generate spike-only data using a Sanger sequencing approach because they lack access to next-generation sequencing or because the former is more cost-effective. Lastly, it is known that coronaviruses can undergo recombination and that the resulting mosaic genomes often contain an alternative spike gene whose ancestry is different to that of other genomic regions. This trend is also observed in those SARS-CoV-2 recombinant genomes that have been discovered so far (Jackson et al. 2021). One consequence of such recombination is that the genetic ancestry of the spike gene may be different to that of the rest of the virus genome.

It is therefore valuable to consider how to assign Pango lineages using SARS-CoV-2 spike-only nucleotide sequences. This question is particularly relevant to the variants of concern (VOC) and variants of interest (VOI), as the specific genomic surveillance of VOC/VOIs is important for outbreak control decisions made by public health agencies. Further, several of the current VOCs contain (and to some extent are defined by) unusually large numbers of distinct mutations in their spike genes that may affect virus antigenicity and molecular diagnostic tools.

The machine learning model in the assignment tool *pangolin* is trained on a data set of genomes that have been designated to Pango lineages using whole genome information (O’Toole et al. 2021). These lineage designations *cannot* be applied directly to a training data set of spike-only sequences because, as we show here, some lineages share identical spike sequences. This would result in the training data containing conflicting or non-identifiable information and would lead to poor model performance. It is therefore important to designate spike-only sequences to a set of taxonomic groups that is appropriate for the information content of spike-only sequences.

Here, we determine the degree to which Pango lineages are distinguishable using spike-only nucleotide sequences, with a focus on current variants of concern. We explore how unique or shared spike mutations observed at differing frequencies may be used to support the classification of spike-only sequences. We introduce the concept of a “lineage set” to represent the range of Pango lineages that are compatible with a given SARS-CoV-2 sequence and we show how each spike-only sequence can be designated to a lineage set using a pre-defined consensus haplotype for each lineage. The resulting lineage set designations provide a platform for the development of Pango lineage assignment tools suitable for spike-only sequences

## Methods

### Data Set Collation and Analysis

We downloaded all SARS-CoV-2 genomes with Pango designations (v1.2.32) available on GISAID on 05^th^ July 2021 (n=458,622). The genomes were aligned against a reference genome for SARS-CoV-2 (Genbank accession ID = NC_045512) using minimap2 v2.17 (Li 2018) and were scaffolded and padded to full-length genome alignments using gofasta (https://github.com/cov-ert/gofasta). Using the Genbank features annotations (accession ID: NC_045512; gene ID: 43740568), we extracted the spike gene nucleotide sequences from the full alignment (genome positions 21,562 to 25,384), resulting in alignments of length 3,822 nucleotides. For each spike nucleotide sequence (SNS) in the alignment, spike ambiguity content was calculated as the proportion of bases present in the spike sequence that was not A, C, G, T, or a gap, and so includes both N and non-N ambiguities in the calculation. Only genomes with complete spike proteins (i.e. no ambiguous bases) were included in subsequent analysis, resulting in 345,356 sequences being included in the analysis (78% of designated sequences). Of the Pango lineages in the dataset, 8 of these lineages had no representative sequences with complete spike protein sequences and were not included in subsequent analysis, leaving 1,296 lineages with complete representative spike proteins.

### Identification of consensus spike haplotypes

To identify consensus spike haplotypes and lineage sets, we first mapped the whole genome sequences of the 345,356 genomes identified above against the Wuhan/WHO4/2020 genome (GISAID accession = *EPI_ISL_406801*) which is a high quality, early, example of lineage A. Using the resulting output alignment file, we identified all non-synonymous mutations, synonymous nucleotide mutations, and insertions and deletions using gofasta (https://github.com/cov-ert/gofasta). We subsequently retained only mutations in the spike protein from these records. Using the Pango sequence designation list (v1.2.32; https://github.com/cov-lineages/pango-designation/blob/master/lineages.csv) we recorded the list of spike mutations observed in each lineage, and the relative frequency of each mutation within each lineage. These lists were then used to calculate a consensus spike haplotype (CSH) for each lineage. Specifically, spike mutations were included in the CSH if they were observed in >X% of the sequences designated to that lineage. Finally, we used the CSHs for each lineage to create lineage sets. These were generated by placing all lineages with the same CSH in the same lineage set. Finally, a unique alphanumeric identifier was generated for each set using the MD5 hash algorithm to allow concise and accurate referencing of the sets.

## Results

### Analysis of SARS-CoV-2 genome and spike gene variation

The extent to which SARS-CoV-2 lineages can be assigned to spike-only nucleotide sequences will depend on the level of genetic variation within and among spike sequences that have been designated to different Pango lineages. Genetic variation within spike can be categorised as (i) non-synonymous single nucleotide changes (i.e. amino acid changes), (ii) synonymous nucleotide changes, and (iii) insertions and deletions (indels).

Figure 1 shows the genomic distribution and frequency of each type of genetic variation (hereafter collectively referred to as mutations) in the GISAID database. The vast majority of amino acids positions in the spike protein (1138 of 1274; 89.2%) exhibit amino acid variation among SARS-CoV-2 genomes. Many non-synonymous changes are rare, found in only one or two genomes, whilst others are shared, or observed in nearly all sequences. Synonymous genetic variation is also abundant across the spike gene and observed at 1302 out of 3822 nucleotide sites (34%). Indel changes, as expected, are considerably rarer (n=158) and concentrated in the N-terminal domain (NTD) of the spike protein. This level of genetic variation suggests that spike-only nucleotide sequences may contain sufficient information to distinguish among many Pango lineages.

**Figure 1.**
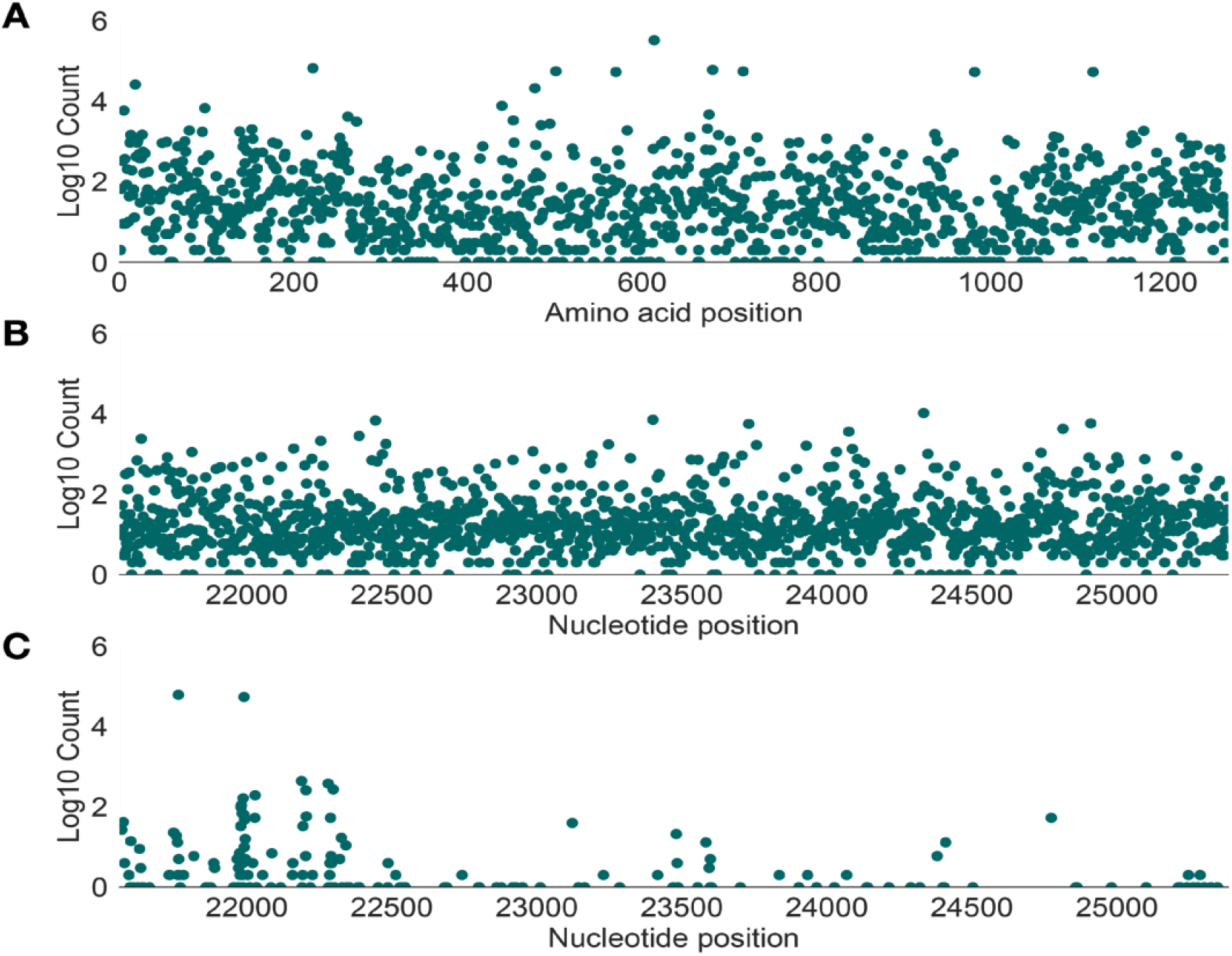
Genetic diversity in the spike protein of all available SARS-CoV-2 genome sequences. The vertical axis of each plot shows the number of designated sequences that exhibit mutations at each amino acid position (log10-scaled). (A) Distribution of non-synonymous variation across amino acid positions of the spike protein (horizontal axis). (B) Distribution of synonymous variation across nucleotide sites in the spike protein (horizontal axis). (C) Distribution insertion and deletion variation (indels) across nucleotide sites in the spike protein (horizontal axis). Each point represents the 5’ nucleotide site at the start of each indel. In each plot, genetic variation is determined by comparison with a lineage A reference sequence (Wuhan/WH04/2020, EPI_ISL_406801).

A similar analysis of genetic variation within the viral spike protein was performed separately for each of the Pango lineages that correspond to the main variants of concern (Figure 2). The frequency of mutations within each lineage were calculated by comparison to an ancestral reference strain. For each lineage, there are a small number of spike mutations that are observed in all, or nearly all, sequences and which form a row along the top of each plot in Figure 2. These mutations comprise the set of spike changes characteristic of each lineage/VOC. The remainder of the observed mutations in each lineage are generally observed at low frequencies, which is the pattern expected by population genetic theory for a rapid-growing population with a single, recent origin (e.g. Slatkin & Hudson 1991). The number of variable spike amino acid sites for each lineage depends on the number of sequences for that lineage. Most amino acid positions in the spike protein of lineage B.1.1.7 (alpha) are genetically variable, although almost all mutations are observed at low frequencies. The number of variable sites for B.1.351 (beta), P.1 (gamma) and B.1.617.2 (delta) are considerably lower, commensurate with the smaller number of sequences available for them with Pango lineage designations (note that many recent VOC genomes deposited on GISAID may have been assigned to a lineage using pangolin but not yet been given a formal lineage designation). As in Figure 1, most indel mutations are observed in the spike N-terminal region of the spike protein.

**Figure 2.**
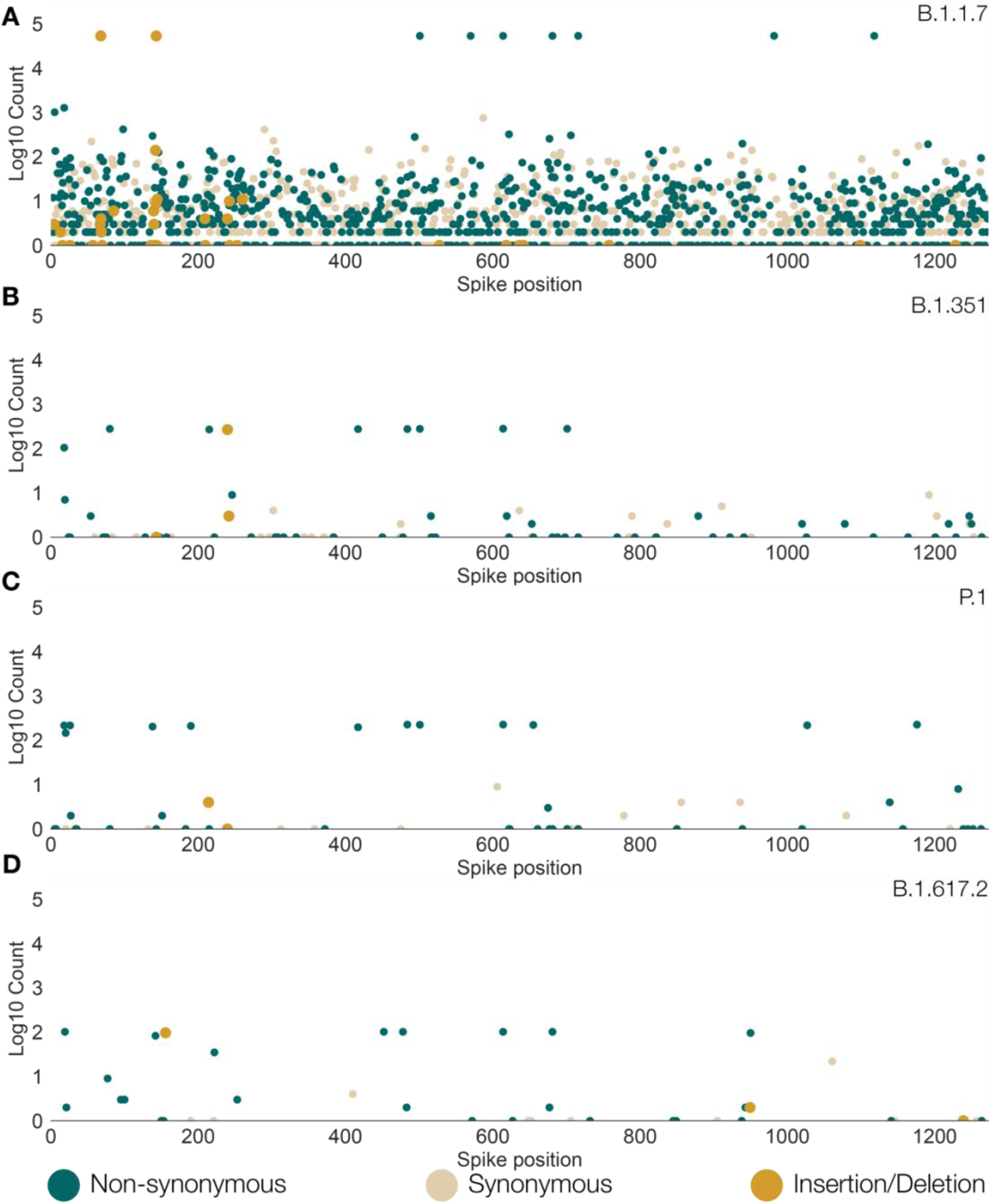
Genetic diversity in the spike protein of SARS-CoV-2 sequences for each of the Pango lineages that correspond to the four main variants of concern. Each coloured dot shows the genomic position of a non-synonymous (green), synonymous (beige), or indel (orange) mutation. The vertical axis shows the number of designated sequences that exhibit non-synonymous variation at each amino acid position (log10-scaled). In each plot, mutations were defined by comparing sequences with a common reference strain, Wuhan/WH04/2020 (EPI_ISL_406801). (A) Lineage B.1.1.7 (alpha). (B) Lineage B.1.351 (beta). (C) Lineage P.1 (gamma). (D) Lineage B.1.617.2 (delta).

The spike nucleotide sequences of designated SARS-CoV-2 genomes available from GISAID (n=345,356) are not all complete (Figure 3). Although by definition such sequences must have <5% of sites with ambiguities across the whole genome, many contain spike sequences that comprise >10% or >20% of ambiguous positions. Overall, 21.9% contain at least one IUPAC nucleotide ambiguity code. The presence of ambiguous sites will hinder our ability to uniquely allocate spike-only sequences to specific Pango lineages, especially as some lineages are genetically closely related. There are a small number of Pango lineages with no complete spike nucleotide sequences (i.e. the spike sequences of all designated genomes within the lineage contain at least one IUPAC ambiguity code). Figure 4 illustrates the completeness of spike sequences belonging to these lineages, which typically contain few genomes. These lineages were not included in the subsequent analyses described below.

**Figure 3.**
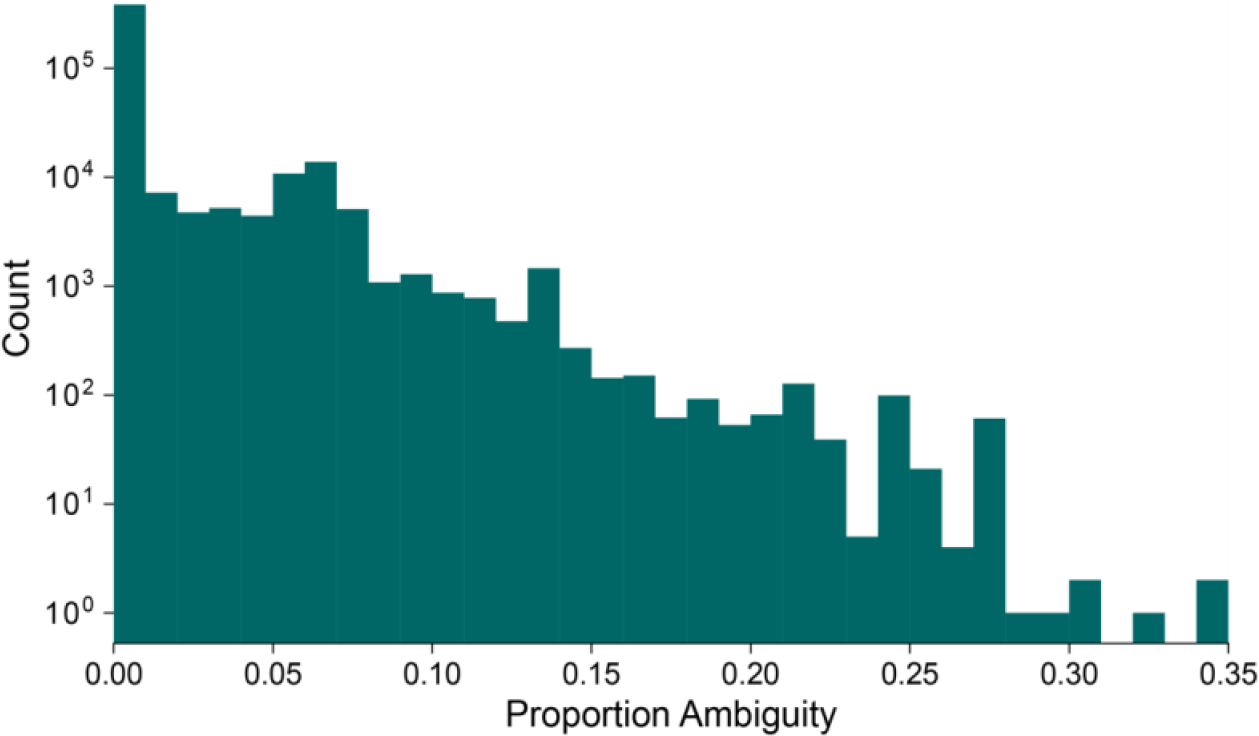
Histogram of the proportion of spike protein nucleotide sites that are ambiguous (i.e. contain at least one IUPAC ambiguity code). The distribution is calculated for all SARS-CoV-2 sequences in GISAID that (i) have been designated to a Pango lineage, and (ii) have N at <5% of sites across the whole genome, excluding UTRs.

**Figure 4.**
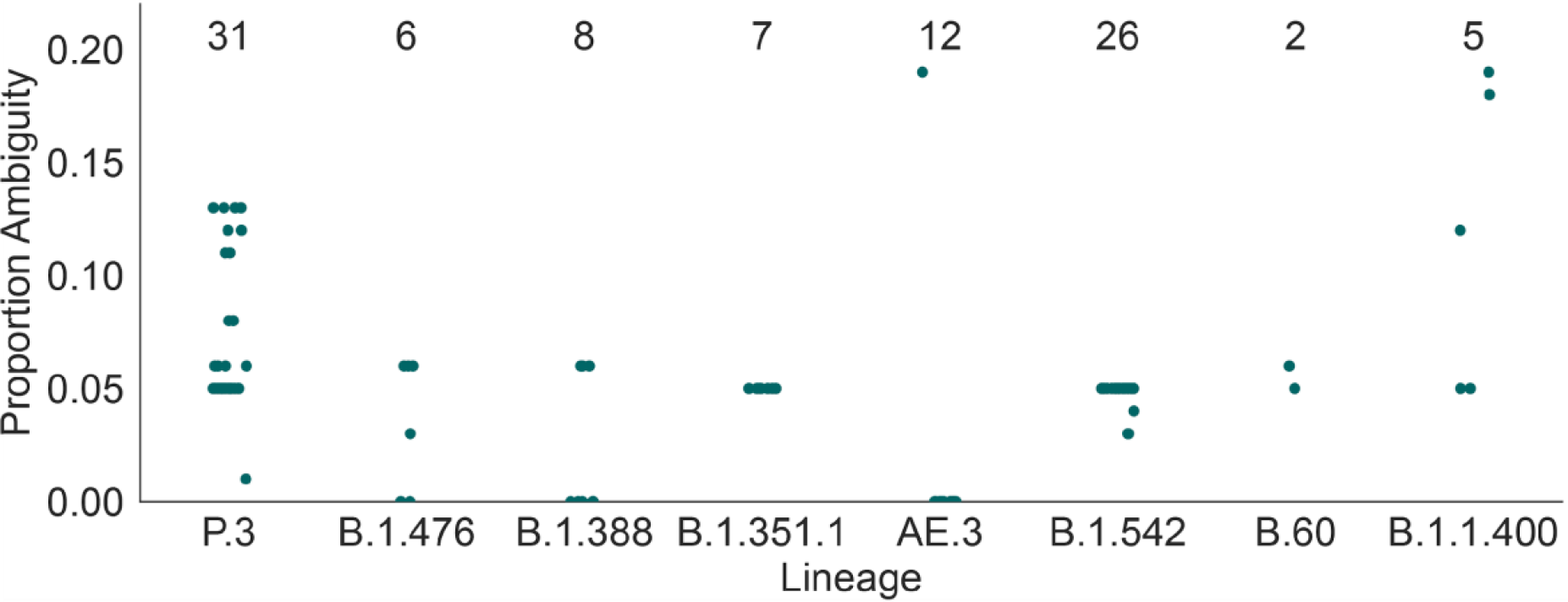
We observed 8 Pango lineages in which every designated sequence contains at least one ambiguity code in its spike nucleotide sequence. The number of sequences designated to these lineages is typically small (range 6-31). For each sequence in these 7 lineages, the y-axis shows the proportion of spike nucleotide sites in that sequence represented by an IUPAC ambiguity code (i.e. not A, C, G, T, or a gap).

### Spike nucleotide sequences

We next examined the degree to which Pango lineages can be accurately distinguished using whole spike nucleotide sequences (SNS). In the Pango lineage designation list (v1.2.32) analysed here (https://github.com/cov-lineages/pango-designation/blob/master/lineages.csv), there are 458,622 genomes with a lineage designation. Among these sequences there are a total of 31,940 different SNSs. Although the number of distinct SNSs is greater than the number of Pango lineages (n=1,304), the same SNS is often found in multiple Pango lineages (Figure 5). Specifically, among the 31,940 SNSs, ∼2,600 are observed in more than one Pango lineage and 29,341 are observed in only one Pango lineage. Importantly, four SNSs are found in >50 Pango lineages and one is found in 658 different lineages (defined as the reference sequence with mutation D614G; Figure 5).

**Figure 5.**
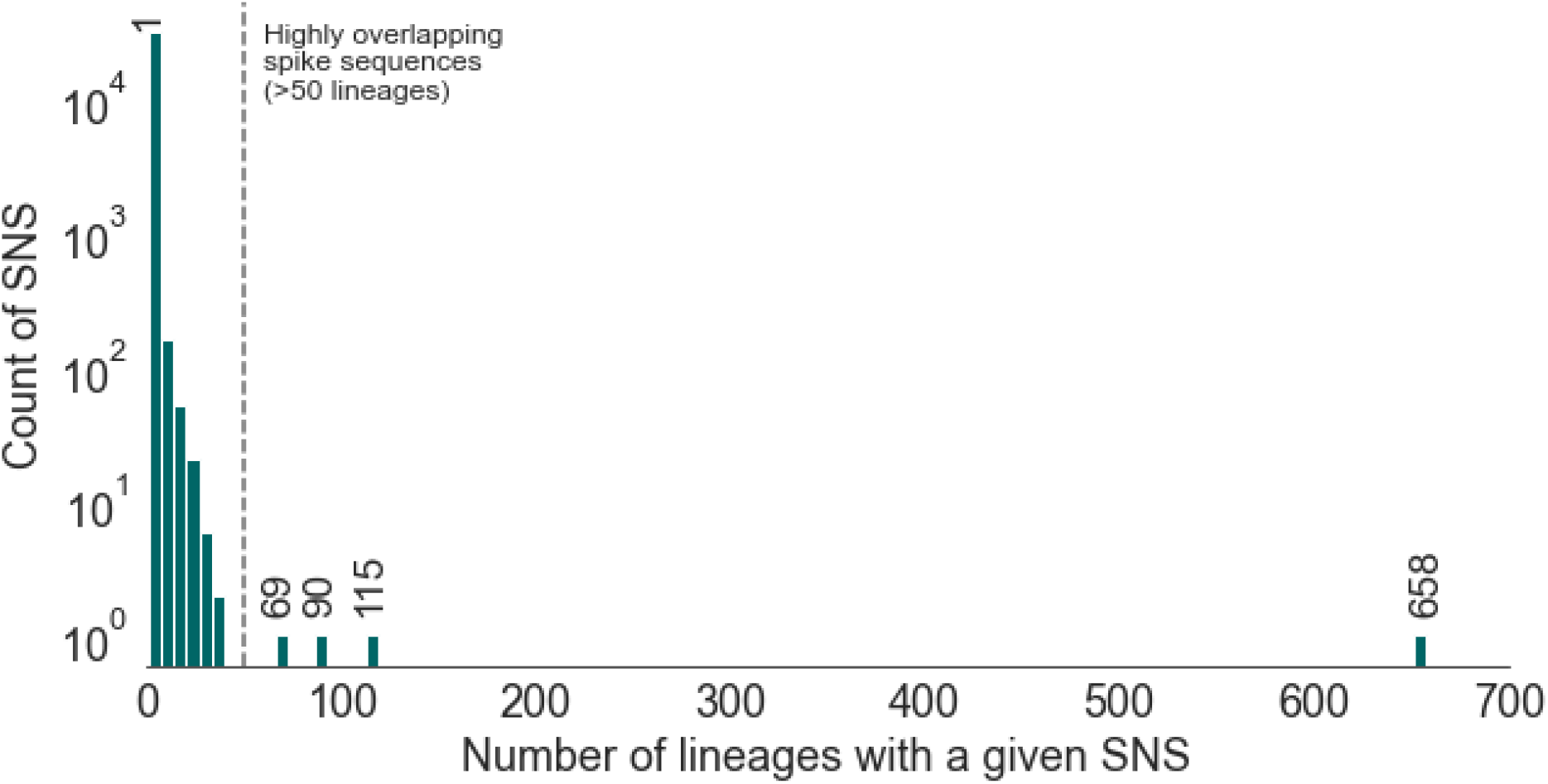
Distribution of the number of lineages that a given SNS is observed in. Most SNS are found in only one lineage. However, a few SNSs can be found in many lineages.

Furthermore, there is high variance in the number of SNSs observed in each Pango lineage (Figure 6). In our data set, the three lineages containing the largest number of SNSs are B.1.1.7, which contains 3,893 SNS, B.1.177 with 2,974, and B.1 with 2,763 (Figure 6). However, the majority of lineages contain very few different SNSs, and >1,000 lineages contain only one SNS. Thus unique spike sequences cannot unambiguously discriminate among all Pango lineages, because some sets of lineages are genetically similar, or differ only at genomic positions outside of the spike gene.

**Figure 6.**
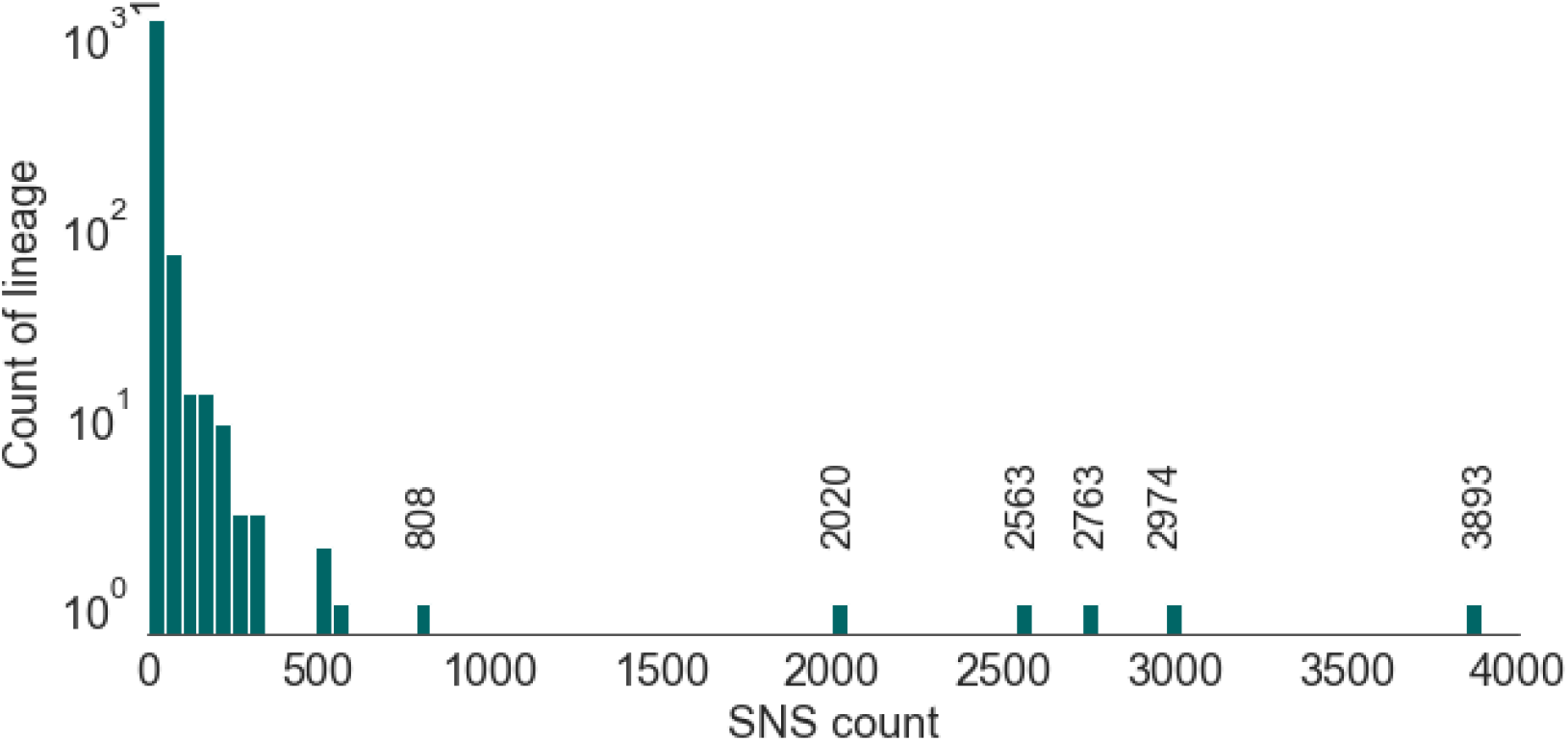
Number of distinct spike nucleotide sequences (SNS) in designated Pango lineages with complete spike sequences. Some of the very large lineages, for instance B.1.1.7 (count=3,893), B.1.177 (count=2,974), and B.1 (count=2,763) have many SNS, whereas the majority of lineages have few.

### Consensus spike haplotypes

The above analysis of SNSs shows that spike-only sequences cannot unambiguously determine the Pango lineage to which a genome belongs, and that some lineages contain large numbers of distinct spike sequences. Further, the number of SNS within a lineage will increase as the lineage diversifies and as new genomes are generated and reported through time. Therefore it will be challenging to maintain a stable and complete list of SNSs for each lineage.

We therefore considered an alternative approach to evaluating the information about Pango lineage in spike-only sequences. Specifically, we analysed the spike sequence diversity within each Pango lineage, then determined which mutations/indels best represent that diversity. This approach is related to the definition of a consensus haplotype for a group of sequences. Given a mutation frequency threshold, X, and a specified lineage Y, we can seek to identify all the mutations/indels that are observed in at least X% of the spike sequences belonging to lineage Y. This set of sites can be then used to define a “X% consensus spike haplotype” for lineage Y. Like SNSs, the consensus spike haplotypes (CSH) will not be unique to a Pango lineage, but will be easier than SNSs to curate, and simpler to interpret because they define a set of common characteristic spike mutations for each lineage. Further, CSH will be more robust to the presence of sequencing errors and low-frequency deleterious mutations.

Many CSHs will be unique to a one Pango lineage, however others will be shared among multiple lineages. We therefore introduce the concept of a “lineage set”, which comprises the set of lineages that share the same CSH. Lineage sets are dependent on the mutation frequency threshold X% used to define the consensus spike haplotype for each lineage. Lineage sets further capture the degree of uncertainty in the process of designating Pango lineages using spike-only sequences.

We apply a conservative mutation frequency threshold of X=95%, i.e. the CSH for a lineage is the set of all mutations observed in 95% or more of the spike sequences in that lineage. We discuss later the problem of choosing a suitable value for X. Using X=95%, we observe a total of 393 CSHs in the data set analysed here (https://github.com/cov-lineages/pango-designation/blob/master/lineages.csv). Among these CSHs, 56 are observed in more than one Pango lineage and 337 are found in only one Pango lineage. Figure 7 shows the size of the lineage sets for the CSHs defined by a mutation frequency threshold of 95%. The most common CSH is shared by 616 Pango lineages and is defined by only one spike non-synonymous mutation, 614G. As expected, the CSHs shared by many lineages are characterised by a comparatively small number of spike mutations. The second-largest lineage set, comprising 75 Pango lineages, contains no mutations and therefore represents spike sequences identical to the ancestral reference sequence. Importantly, the four Pango lineages that correspond to the main VOCs (B.1.1.7, B.1.351, P.1, B.1.617.2) each have a unique CSH that is not shared by other lineages, and therefore these lineages can be classified well using spike-only nucleotide sequences.

**Figure 7.**
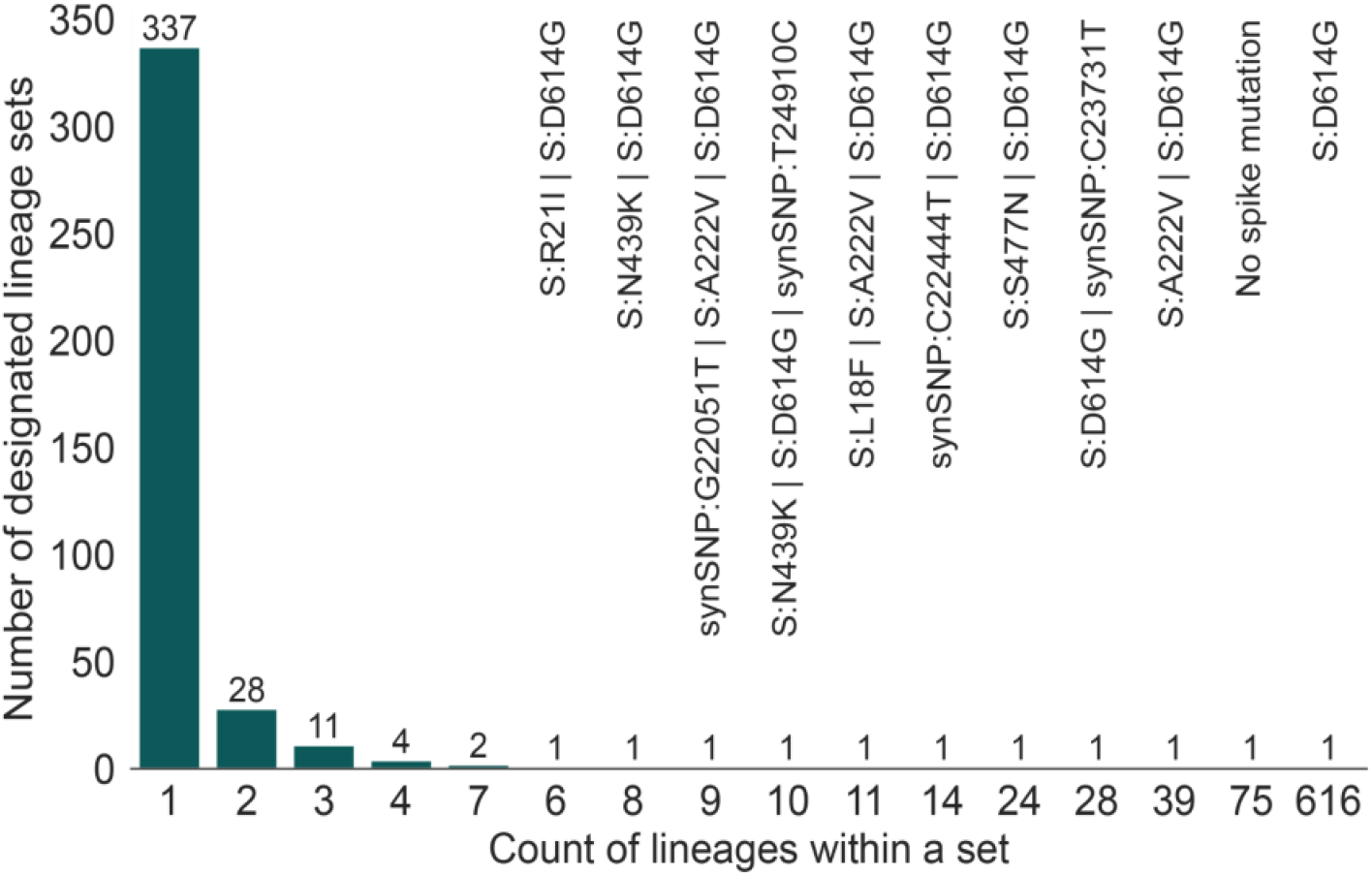
Distribution of the number of Pango lineages in each lineage set, using a mutation frequency threshold of X=95%. 337 sets contain only a single lineage and so are uniquely distinguishable by their consensus spike haplotype. 28 sets contain two Pango lineages, and so on.

### Nomenclature for lineage sets

A complete list of the 393 lineage sets obtained using a 95% mutation frequency threshold is available online at https://github.com/aineniamh/hedgehog/blob/main/hedgehog/data/set_names.95.csv. For some lineage sets, the number of lineages within the set can be large, hence a list of the lineages is not a convenient way to name or refer to these lineage sets. Therefore to facilitate communication, we use a simple hashing algorithm to convert the list of lineages in each lineage set into a unique six-character alphanumeric code (see Methods). Using these names, lineage sets can be conveniently compared between data sets and analyses. Details of the six-character names of the 393 lineage sets are provided online at the URL above. For example, the name of the large lineage set defined by a X=95% CSH comprising only mutation 614G is named “d03608”. The lineage set that contains only lineage B.1.1.7 (alpha) is named “2430b9” and corresponds to the X=95% CSH that contains the following spike mutations: del:21765, del:21991, N501Y, 570D, 614G, 681H, 716I, 982A, and 1118H

### Choice of mutation frequency threshold

In the analysis above we used a mutation frequency threshold value of X=95% to define CSHs. Values of X closer to 100% are unlikely to be useful because mutations that are highly characteristic of a lineage are not be found in all designated sequences in that lineage, because recurrent mutation, back mutation, or sequencing error may lead to the loss of a spike mutation in a small proportion of sequences in a lineage. This phenomenon is illustrated in Figure 8, which shows how the number of mutations in the CSH for a lineage declines as X increases. The number of mutations drops sharply as X approaches 100%, especially for the four main VOC lineages, which all contain a large number of spike mutations in their CSHs. The presence of many low-frequency spike mutations is evident in the rapid decline in the number of mutations as X increases from 0 to 10%.

**Figure 8:**
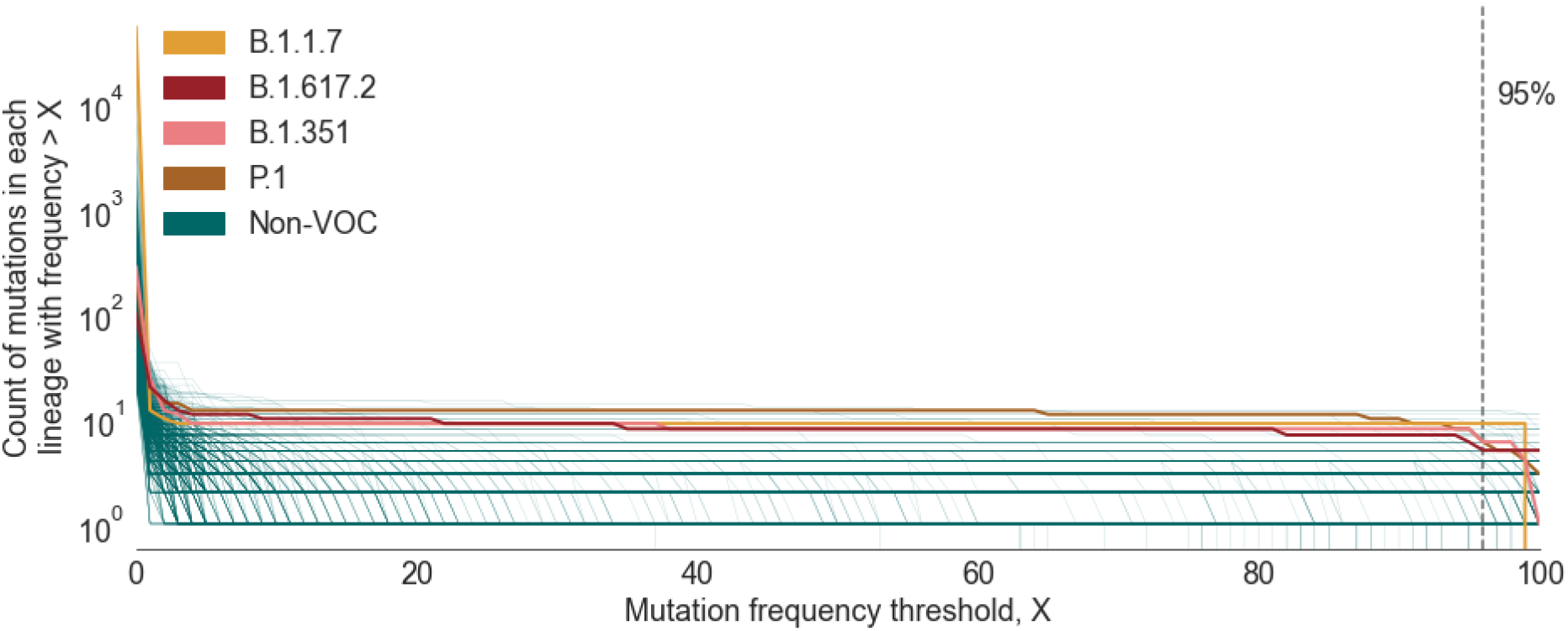
Plot showing the number of mutations in the CSH for a given lineage as a function of the mutation frequency threshold (X) used to define the CSH. Lineages shown are those that contain >20 designated sequences with complete spike nucleotide sequences and >5 spike mutations. The lineages that correspond to the four main VOCs are coloured individually, whilst all other included lineages are shown in green.

Mutations may arise within a lineage as it grows and diversifies and these will be observed in only a fraction of sequences designated to that lineage. Consequently, as X decreases, the probability increases that a given spike sequence (which has been designated to lineage Y, say) does not contain all of the mutations in the CSH for lineage Y and cannot be designated to any lineage set. Therefore the choice of X represents a trade-off between the specificity and sensitivity of designating spike-only sequences using lineage sets. This trade off is shown in Figure 9. If X is too low, many sequences can’t be given a lineage set designation (Figure 9a), whilst if X is too high, the average number of lineages in a lineage set is large (Figure 9b), which will reduce specificity of lineage assignment. The mutation frequency threshold of X=95% is suitable if we wish to designate a high proportion of spike-only sequences. A lower threshold value could be chosen if specificity is to be prioritised.

**Figure 9:**
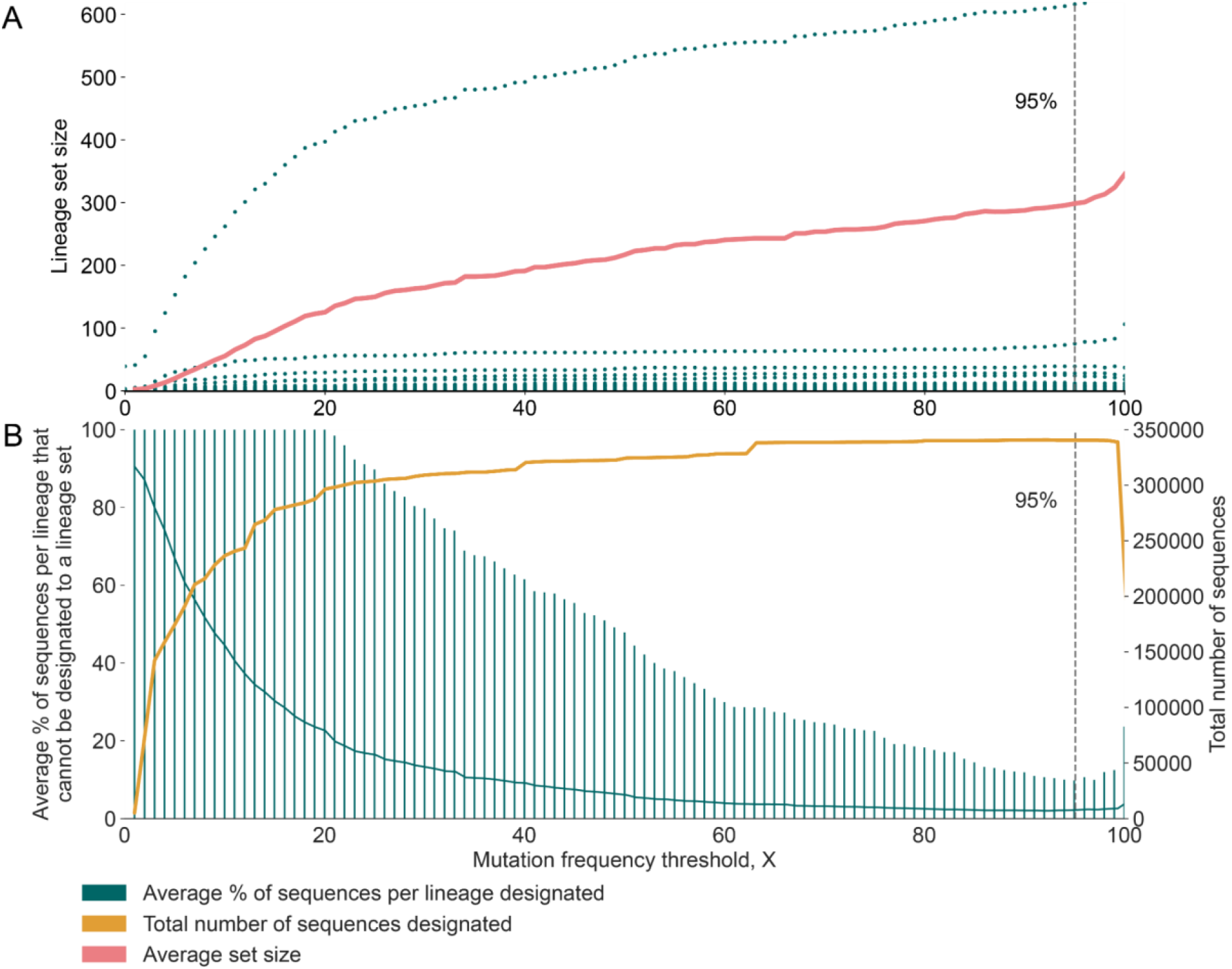
Plot showing the trade-off arising from the choice of mutation frequency threshold, X. (A) The mean number of lineages in each lineage set increases as X increases from 0 to 100% (pink). The mean is calculated across all spike-only sequences that can be designated to a lineage set. The mean is dominated by one very large lineage set; green dots show the actual set sizes for all lineage sets. (B) The total number of sequences that can be designated to a lineage set increases as X increases from 0 to 100% (orange). The mean percentage of sequences per lineage that cannot be designated to a lineage set is also shown in order to normalise the effect of lineage size (green). Error bars represent 95 percentiles of this distribution.

## Discussion

Spike-only SARS-CoV-2 nucleotide sequences constitute an important source of genetic information about the virus, which are of particular relevance to host receptor binding, humoral immune responses, and current variants of concern. Here we explore how such sequences can be classified within the Pango lineage nomenclature, which was developed initially for complete virus genome sequences. Despite the significant amount of genetic variation among the spike genes of available SARS-CoV-2 genomes, we do not find a one-to-one mapping between spike nucleotide sequences/haplotypes and Pango lineages. Although most spike nucleotide sequences (SNS) are observed in only one Pango lineage, many SNS are observed in multiple lineages, and one sequence is found in >600 different lineages. The same is true for consensus spike haplotypes (CSH), which are defined for each lineage by the set of mutations that are observed above frequency X% in that lineage.

We therefore developed a system by which spike-only sequences can be designated to a “lineage set” that contains all of the Pango lineages consistent with the CSH of that sequence. In many instances, the lineage set will contain only one Pango lineage, but in others it may contain tens or hundreds of lineages. These lineage sets reflect our uncertainty in Pango lineage designation from partial sequences, and this interpretation extends readily to sequences that contain ambiguous or missing bases. We note that lineage sets could in theory be applied to SARS-CoV-2 sub-genomic regions other than spike.

In order to ensure that lineage set designations are robust to common minor sequence variants within each lineage, our results indicate that lineage sets should be designated using CSHs that are defined with a mutation frequency threshold (X) lower than 100% (Figure 9b). Training a custom pangoLEARN model on spike-only sequences that have been designated to such lineage sets will circumvent the conflicting information (and consequent errors in assignment) that would arise if the model were to be trained on spike-only sequences designated to Pango lineages. Training the pangoLEARN model on lineage set designations avoids the issue of missing data outside of the spike gene and will allow all spike sequence variation (not just the mutations that contribute to the CSH) to inform the assignment. Once a reference set of spike sequences has been designated to lineage sets using the CSH approach outlined here, it is relatively straightforward to apply these designations to pangolin (O’Toole et al 2021) and consequently assign spike-only sequences to the lineage sets. For convenience, we have written a python-based script that can output lineage set assignments based on a custom pangoLEARN model trained on spike sequences. This tool is freely-available at https://github.com/aineniamh/hedgehog.

SARS-CoV-2 genomes have diversified and diverged as the pandemic has unfolded, and large numbers of mutations in the spike gene are reported in the VOCs that have emerged since late 2020. We expect spike sequences to continue to diverge, so that in the future we expect that it will become easier to assign Pango lineage using spike-only sequences. Virus genomes sampled at the start of the pandemic were genetically similar to each other, hence the SNS and CSH shared among many lineages tend to be those genetic types that were present early in the pandemic. If we wish to reduce the size of the large lineage sets, then it will be necessary to use information about virus genomes other than their spike nucleotide sequences. In theory, the sequence sampling date and/or location could be used; if certain lineages are known to have been highly prevalent or absent from a region at different times, then that information could be used, in a Bayesian frameowrk, to inform the assignment inference. However, such an approach might be suitable only for those regions or countries that have undertaken consistently high levels of unbiased population-level genomic sampling and surveillance.

## Data availability

All software and source code is available on GitHub at https://github.com/aineniamh/hedgehog.

